# Chimera States among Synchronous Fireflies

**DOI:** 10.1101/2022.05.12.491720

**Authors:** Raphaël Sarfati, Orit Peleg

## Abstract

Systems of oscillators, whether animate or inanimate, often converge to a state of collective synchrony, when sufficiently interconnected. Twenty years ago, the mathematical study of models of coupled oscillators revealed the possibility for complex phases which exhibit the coexistence of synchronous and asynchronous clusters, since then referred to as “chimera states”. Beyond their recurrence in theoretical models, chimera states have been observed in specifically-designed, non-biological experimental conditions, yet their emergence in nature has remained elusive. Here, we report robust evidence for the occurrence of chimera states in a celebrated realization of natural synchrony: fireflies. In video recordings of collective displays of *Photuris frontalis* fireflies, we observe, within a single swarm, the spontaneous emergence of different groups flashing with the same periodicity but with a constant time delay between them. From the three-dimensional reconstruction of the swarm, we demonstrate that these states are stable over time and spatially intertwined, but find no evidence of enhanced correlations in their spatial dynamics. We discuss the implications of these findings on the synergy between mathematical models and firefly collective behavior.

## 1 Introduction

Complex systems consisting of entities with internal periodicity often produce synchrony. This has been observed, demonstrated and characterized across (spatial and temporal) scales and ensembles [1, 2], animate or inanimate, from planetary orbits, to ecosystems [3], animal collectives [4, 5], cell tissues (cardiac or neuronal), and down to electronic structures. The underlying reason for such ubiquity is that interacting oscillators, even weakly coupled, tend to adjust their individual frequencies and drift towards a common phase. This is what the mathematical analysis of simple models of coupled oscillators has uncovered [1, 6, 7]. However, if such models converge to synchrony for a wide range of coupling schemes, the modalities of the resulting synchronous patterns can be quite complex, and notably include different phases [8]. Among them, recent research has focused on phases that exhibit a coexistence of synchronous and asynchronous dynamics [9], where constituting agents separate into different clusters aligned on different tempos. Such phases have been named “chimera states” in reference to the Homeric hybrid creature made of parts of disparate animals [10]. In a chimera state, coexisting subpopulations can either be synchronous and asynchronous, or mutually synchronized on different tempos. While abundant in mathematical models, chimeras remain usually rare in the real world. Certain chimera states have been observed in carefully-designed experimental systems, yet they remain wildly elusive in nature, and biological settings in particular [11].

In this paper, we present evidence for the occurrence of chimera states in natural swarms of *Photuris frontalis* fireflies. These fireflies are one of few species known for their precise and continuous synchrony [12, 13], with coherent displays which can span several tens of meters. Synchronous fireflies, wherein congregating males flash in unison possibly to optimize signal communication with grounded females [14], have long been considered a picturesque paragon of natural synchrony and an inspiration for theoretical developments. Yet, until recently, little was known about the details of their collective dynamics, in particular their spatiotemporal patterns [15]. Based on high-resolution stereoscopic video-recordings, we demonstrate the existence and persistence of synchronized chimera states within *P. frontalis* swarms. We characterize their spatial distribution and movement, and find that chimeras appear spatially intertwined while slightly clustered without enhanced correlations in their displacement, and generally stable in their phase difference. We conclude by discussing the theoretical conditions for the emergence of such chimeras, possible implications about the structure of firefly interactions, and how the natural system might further inform future mathematical models.

## 2 Results

In late May in Congaree National Park, *Photuris frontalis* displays ardently across a forest of loblolly pines spreading along the convergence line between a bluff and the Congaree River floodplain [16]. While the swarm stretches across hundreds of meters, most fireflies tend to coalesce into localized leks hovering above smaller parcels. Many fireflies were observed to swarm in an area at the outskirt of the pine forest forming roughly a quadrilateral of side-length ~40m (Fig. 1A). Two cameras were recording at 60 frames-per-second towards the center and above a clear ground (Fig. 1B). From stereo-recording, a portion of the swarm could be reconstructed in 3D, corresponding to a cone of aperture ~35° and length ~40m (Fig. 1A). Recording started at dusk every night, just when the first flashes could be observed, and continued for about 150min.

**Figure 1:**
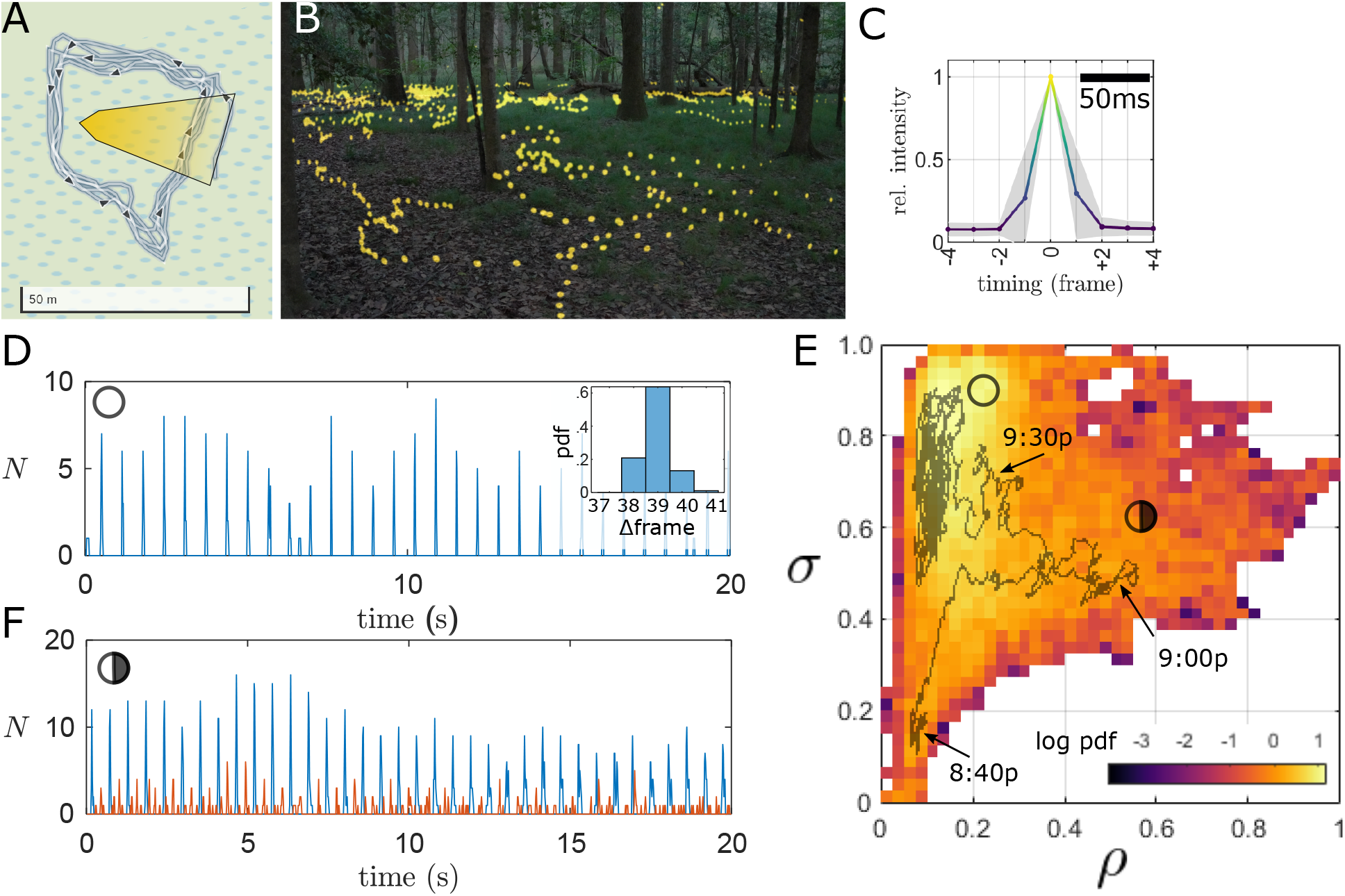
Recording of collective display. (A) Approximate contour of swarming area and overlap of both cameras’ fields of view (yellow cone). (B) Composite image showing the swarming environment as seen from left camera and firefly flashes over 20s. (C) Intensity profile of a single flash over a 150ms time window (average from 100 separate flashes, normalized by maximum value; standard deviation in grey). After thresholding, each flash is typically detected in a single frame due to narrow profile. (D) Time series of the number of flashes per frame, *N*, revealing a chorus of synchronous flashes occurring with great regularity every 0.65s. Inset: distribution of peak-to-peak intervals. (E) Distribution of synchrony *σ* vs density *ρ* (data from every night May 20 - May 30, 140-180min), and averaged trajectory (May 29). Approximate corresponding times (EST) indicating with arrows. The circular markers correspond to the approximate locations of the time series in panels D and F. (F) Time series showing two concurrent and interlaced synchronized groups: the chorus (blue) and dissonant characters (orange). From May 23 around 8:55pm.

*P. frontalis* fireflies produce brief flashes lasting 20ms to 30ms (Fig. 1C), owning them the nickname of “Snappies” [17]. After image binarization by intensity thresholding, each flash is generally detected in a single frame of the movie, sometimes two. They are also known for their precise, collectively continuous synchrony [12]. Indeed, the time series of the number N of flashes per frame reveals a *chorus* of sharp spikes of up to 20 concurrent flashes repeated with great regularity (Fig. 1D). Here, the duration between these collective flashes is narrowly distributed around 39 ± 1 frames (Fig. 1D Inset), or 0.65 ± 0.02s, although this period is strongly inversely correlated with temperature [13, 17], and hence varies over time (typically between 0.5s and 1s, Fig. S1).

Over the course of the evening, the collective flashing density *ρ*, quantified as the sliding average of the number of flashes per frame, increases rapidly for 15min, then decreases slowly and eventually plateaus from 9:30pm until after 11pm (Fig. S2. In prior studies of synchronous fireflies, a high degree of synchronization is reached only at high flashing density [18, 19]. However, we remark that this is not the case here. We define the degree of synchronization *σ* ∈ [0, 1] as the number of flashes in the chorus over the total number of flashes, within 1min time intervals (see Methods). The distribution of *σ* versus *ρ* in Fig. 1E shows a noticeable pattern, as the collective display transitions between three different states. Early in the evening, the density is low and flashes tend to be uncorrelated (*σ* ≃ 0.2), while later the same low density produces a highly synchronized display (*σ* ≃ 0.6 - 1). The difference might be due to residual ambient light at earlier times, which can impede visual interactions (see SM). Surprisingly, when *ρ* is at its peak (*ρ* ≥ 0.35), the group is only partially synchronized, with *σ* ≃ 0.5. We remark from the time series that while a majority of flashes concur with the chorus (*χ_0_*), a significant portion occur off-beat, often mutually in unison (Fig. 1F). These nonconformist *characters* (*χ_a_*) form an eclectic ensemble consisting of both independent flashers and sporadic synchronized clusters. In other words, they appear to form a natural occurrence of a chimera state. We resolve to further investigate this type of anecdotal realizations of ambivalent states, by looking at 11 nights of data (May 20-30,2021) in intervals where *ρ* > 0.35.

To study these two interlaced populations, we identify the chorus as those in the immediate temporal vicinity of local maxima in *N* (see Methods) and set apart the characters as all other flashes. Frequency spectra reveal that both *χ_0_* and *χ_a_* exhibit the same periodicity (Fig. 2A). This suggests that the collective dynamics consists of distinct groups of similar oscillators with different phases. While “chimera states” broadly designates the coexistence of synchrony and asynchrony [11], we find that a significant proportion of characters are also synchronized between them. The distribution of *N*(*χ_a_*) shows that about 25% of character flashes occur concurrently with at least one other within the camera’s field of view (Fig. 2B). These statistical signatures suggest the existence of chimera states consisting of at least two several mutually exclusive synchronous clusters.

**Figure 2:**
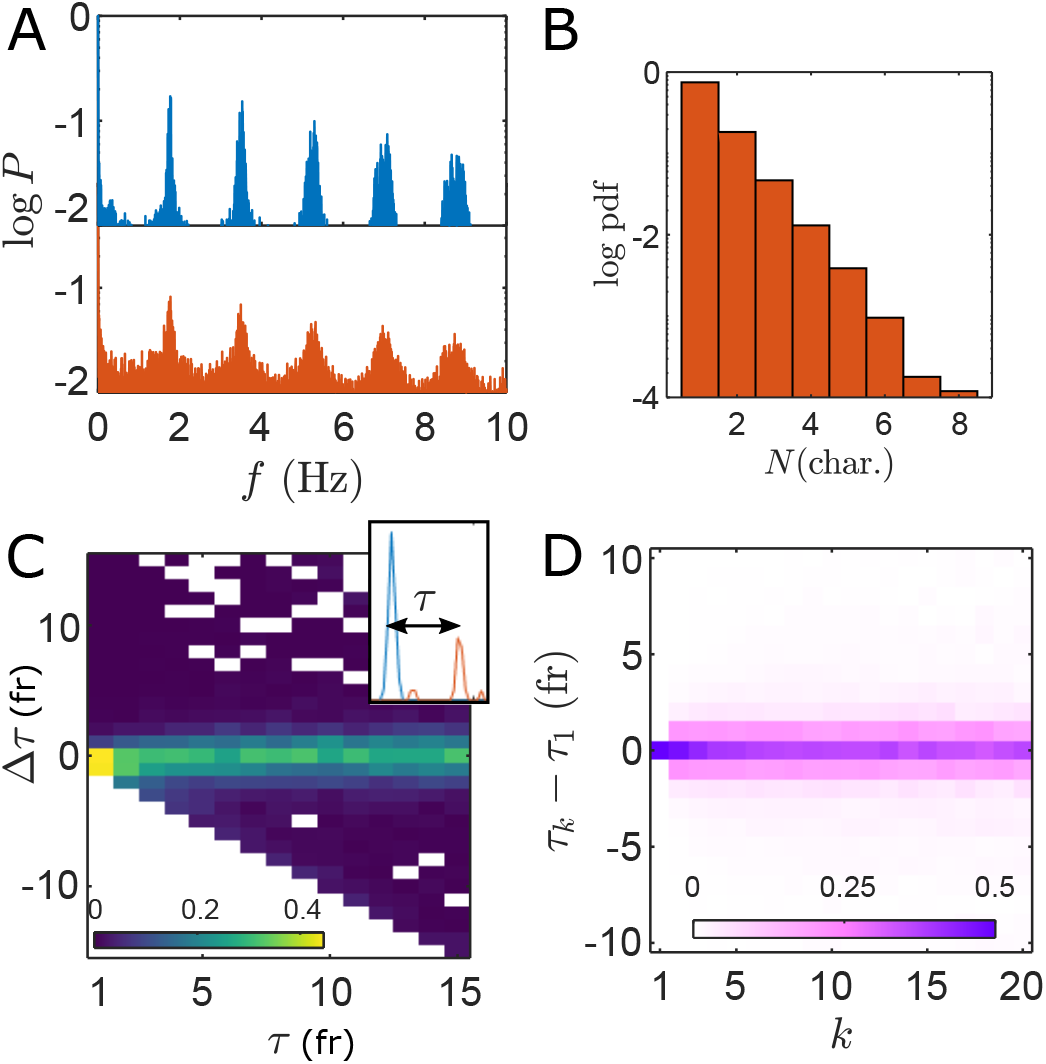
Periodicity and stability of characters. (A) The power spectra P shows that both the chorus and characters have the same periodicity (May 23, over 1 hr). (B) Distribution of the number of concurrent character flashes. About 25% of them have *N* ≥ 2. (C) Using 3D reconstruction, individual trajectories can be tracked. Each flash can be assigned a delay *τ_i_* with respect to the nearest chorus beat (Inset), and *τ*_*i*+1_ for the following flash within the same trajectory. The distribution of flash difference Δ*τ* = *τ*_*i*+1_ *τ_i_* shows that most trajectories maintain the same delay compared to the chorus, independently of what that delay is. (D) Change in time delay for the k^th^ flash compared to the first flash in the trajectory remain narrowly distributed around 0 ± 1 frame even after 20 successive flashes in the trajectory.

Are these chimera states stable? In other words, do off-phase synchronous clusters tend to drift back towards the chorus’ tempo over time or rather maintain their phase difference? To investigate, we look at the temporal evolution of character time delays, defined as the time *τ* between a flash and the closest preceding or following chorus beat. Flash localization in 3D combined with a distance-based linking algorithm allows to track individuals for the duration of their wanderings within the field of view. For each trajectory, we evaluate the difference in time delay Δ*τ* = *τ*_*k*+1_ − *τ*_*k*_ between flash k and succeeding flash *k* + 1. For all initial *τ*, the distribution is sharply distributed around Δ*τ* = 0 (Fig. 2C), indicating that most characters maintain their phase difference with the chorus between flashes. This remains true over several successive flashes, as shown in Fig. 2D. This is strong evidence for chimera persistence.

Having established the existence and temporal stability of chimera states, we characterize their spatial self-organization. Characters and chorus appear generally intertwined, occupying the same space without clear partitioning (Fig. 3A). To demonstrate the absence of topological structure, we consider instances when at least three synchronized characters populate the field of view. In about 20% of these occurrences, at least one member of the chorus is located within the convex hull defined by the group of characters (Fig. 3B). This demonstrates the absence of separation more accurately than looking, for example, at nearest-neighbors, which may be situated outside of the field of view. However, despite the spatial blending of *χ_0_* and *χ_a_*, we find weak signature of group aggregation. For synchronous groups of characters, the distribution of pairwise distances shows that characters are closer together than members of the chorus. The largest distances happen between synchronized characters and the embedding chorus.

**Figure 3:**
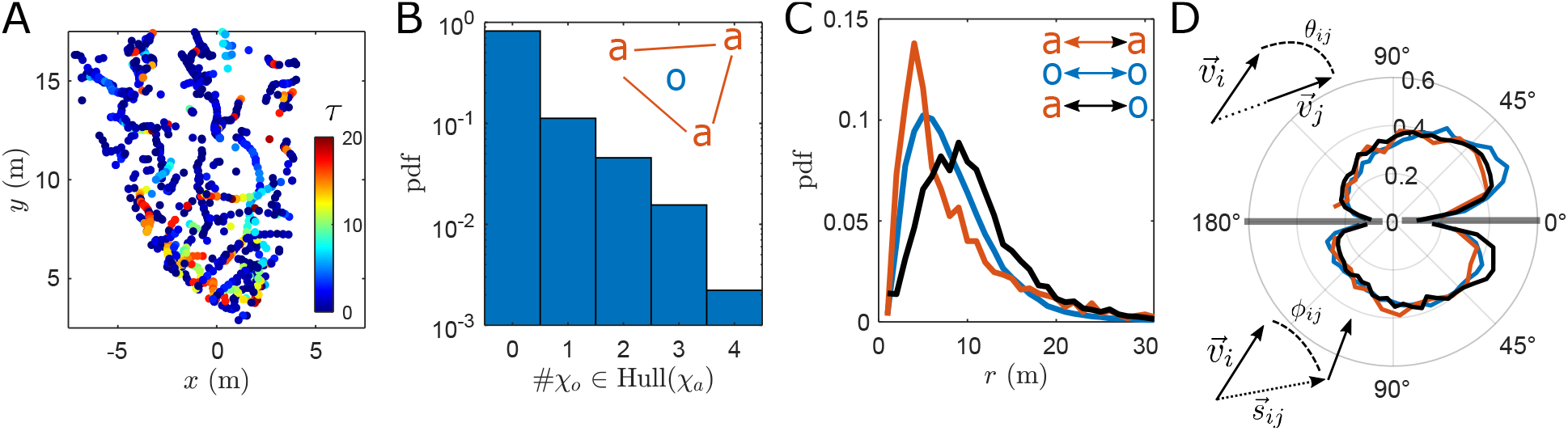
Spatial organization and dynamics of chimeras. (A) Reconstructed top-down view of the swarm in a chimera state (2min interval). Each dot represents a flash, with colors indicating the relative delay *τ* (in frames). Characters appear spatially intertwined with the chorus (dark blue). (B) Distribution of the number of chorus flashes occurring inside a convex hull of synchronized characters (when at least three). The proportion of #*χ_o_* ≥ 1 is about 20%. (C) Distribution of pairwise distances, among the chorus (blue), characters (orange), and between chorus and characters (black). (D) Top: distribution of angles θ between two simultaneous flashers’ instantaneous velocities, *θ* = arccos 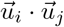. Bottom: distribution of angles ϕ between the velocity of flasher i and its direction towards a simultaneous flasher 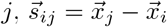. Same colors as in (C).

Finally, we consider firefly kinematics and examine whether separate synchronous groups exhibit correlations in their movement. Due to the geometry of the lek and the positioning of our cameras (Fig. 1A), there appears to be a systematic centripetal bias in firefly displacement which is not merely an artifact of the imaging setup (Fig. S3). Therefore, individual velocities 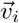 are inherently correlated, but we investigate whether they tend to be more correlated amongst individuals of the same subpopulation. We first look at alignment terms 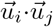 where 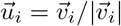. We find no significant difference in the distribution of alignment terms among the chorus or the characters compared to the baseline across population (Fig. 3D). Similarly, considering 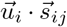, where 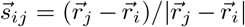, we find no preferential attraction towards individuals of the same subpopulation. These findings, although perhaps surprising, further suggest that spatial and temporal dynamics in natural chimeras need not be correlated.

## 3 Discussion

Twenty years after chimera states were first noticed by Kuramoto and Battogtokh [9], we unearth the existence of comparable states of coexistence between phase-mismatched synchronous clusters in swarms of *P. frontalis* fireflies. Besides the spatially-divided phenomenon of unihemispheric sleep [20, 21, 22], this may be one of the only known occurrences of chimera states in natural or animate systems. Unlike prior experimental realizations of chimeras in systems designed specifically for that purpose [23, 24, 25], firefly chimeras are spontaneous and self-organized. They presumably emerge thanks to a particular type of interactions between individuals and the underlying topology of their connectivity. Using high-resolution, three-dimensional reconstructions of the swarm, we show that these chimeras are persistent and that different synchronous groups are spatially intertwined, although slightly clustered. In the current state, our investigation is somewhat impeded by the fact that only a portion of the swarm can be observed, due to its spread, with fireflies constantly entering and exiting the field of view. Multi-camera systems may be able to improve on this issue and reconstruct the entire lek.

What might the presence of chimera states in firefly swarms reveal about the network of their interactions? Initially, chimeras were believe to only emerge in systems with nonlocal interactions [22]. More recently, they have been discovered in other connectivity structures, namely global or local coupling [26]. All-to-all coupling, where each individual interacts with all others, is evidently not a reasonable assumption in a sprawling group of fireflies with limited field of view. Purely local interactions, where each entity interacts with strictly nearest-neighbors, typically require convoluted, delayed and nonlinear types of interactions to result in fragmentation [27]. Nonlocal interactions, designating weaker or less likely coupling at increasing distances, are the most adequate to produce chimeras [11]. They also reflect a probable social structure in firefly swarms, as supported by simple considerations regarding the extent of line-of-sight interactions as well as prior results in a different synchronous species, *Photinus carolinus* [18].

Reciprocally, spatiotemporal patterns of real-world chimeras may be able to inform and instigate new theoretical paradigms. In particular, current chimera-generating models traditionally involve continuous coupling and a static network of interactions, two assumptions which are evidently at odds with the reality of firefly swarms. Even more perplexing, perhaps, fireflies presumably use cognition in their interactions with each other, a process significantly more complex than the typical functional relations that link abstract oscillators. Natural chimeras, while certainly not malicious, may have *de facto* opened up a Pandora’s box of intriguing new problems for mathematicians to consider.

## 4 Methods

### Field recordings

Field experiments took place at Congaree National Park, South Carolina, USA between May 15 and May 30, 2021 (permit #CONG-2021-SCI-0002). As previously described [18], we used two Sony α7R4 cameras mounted with a wide-angle objective and recording at maximum ISO (32,000) at 60 frames per second. The cameras were about 4m apart, situated and oriented in the same way every night to a very good approximation. Recording started at 8:38pm every night.

### Movie processing

Each night, relative camera pose was estimated from 10 to 15 pairs of images of a checkerboard poster (25cm squares) using Matlab’s stereoCameraCalibrator interface which returned the fundamental matrix for the bifocal system. Frame matching was obtained from the cross-correlation of *N* time series. Flashes were detected in each frame by intensity thresholding after adaptive background subtraction. Flash planar coordinates were then triangulated using the prior estimation of the fundamental matrix.

### Chorus flashes

The times of the swarm’s main beat were computed from the local maxima in the time series. To account for flash persistence, the chorus was defined as all flashes mutually overlapping with beat time (*i.e.*, connected component). The characters were defined as all other flashes.

### Trajectories

Thanks to the low density of active fireflies and the short displacements between flashes from a same individual, trajectories were simply obtained by a distance-based linkage method. Individual “instantaneous” velocities 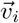 are simply defined as the displacement between two successive flashes within the same trajectory, divided by the time interval between them: 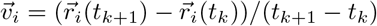.

## 5 Acknowledgements

We thank Congaree National Park and the National Park Service for allowing our research, and David Shelley in particular for guidance in the field. We acknowledge Larry Buschman, Lynn Faust, Julie Hayes, Owen Martin, Andrew Moiseff, Claudia Santiago, Paul Shaw and Mac Stone for precious insights.

## 7 Author Contributions

R.S. conducted experiments, analyzed data and wrote the paper. All authors designed the research and revised the final manuscript.

## 9 Data Availability Statement

Spatiotemporal reconstructions of collective displays are made available as part of the Supplemental Material.

## Supplementary Information to

### 1 Period dependence on temperature

As often noted previously, air temperature has a strong influence on the flashing period of fireflies in general [1] and *Photuris frontalis* in particular [2]. We indeed observe this dependence in our data, with collective flashing periods spanning a range between 0.55s and 0.95s for temperatures between 20C and 27C. The fit to about 30 data points (not shown) in presented in Fig. S1. Collective periods were measured from differences between successive chorus flashing times, and temperature was recorded using a Kestrel 5000 Weather Meter logger.

**Figure S1:**
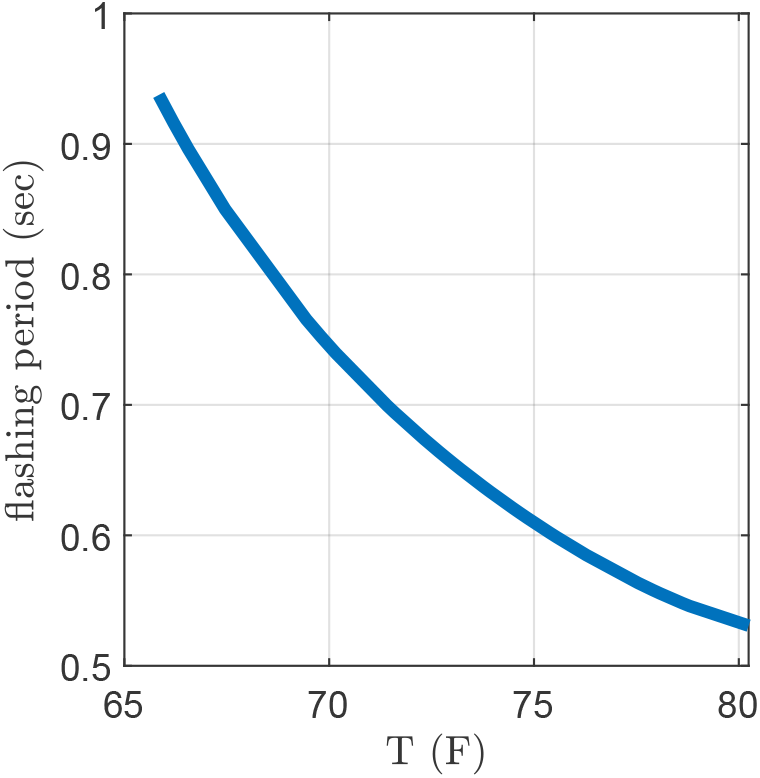
Dependence of collective flashing period on temperature (May 16-May30, 2021).

### 2 Nightly firefly activity

Every night, firefly activity presents a similar pattern, as described in the Main Text. In Fig. S2, we report a typical density profile. In parallel, we plot a measure of residual ambient light after sunset. It falls rapidly until about 9pm, after which obscurity is optimal.

**Figure S2:**
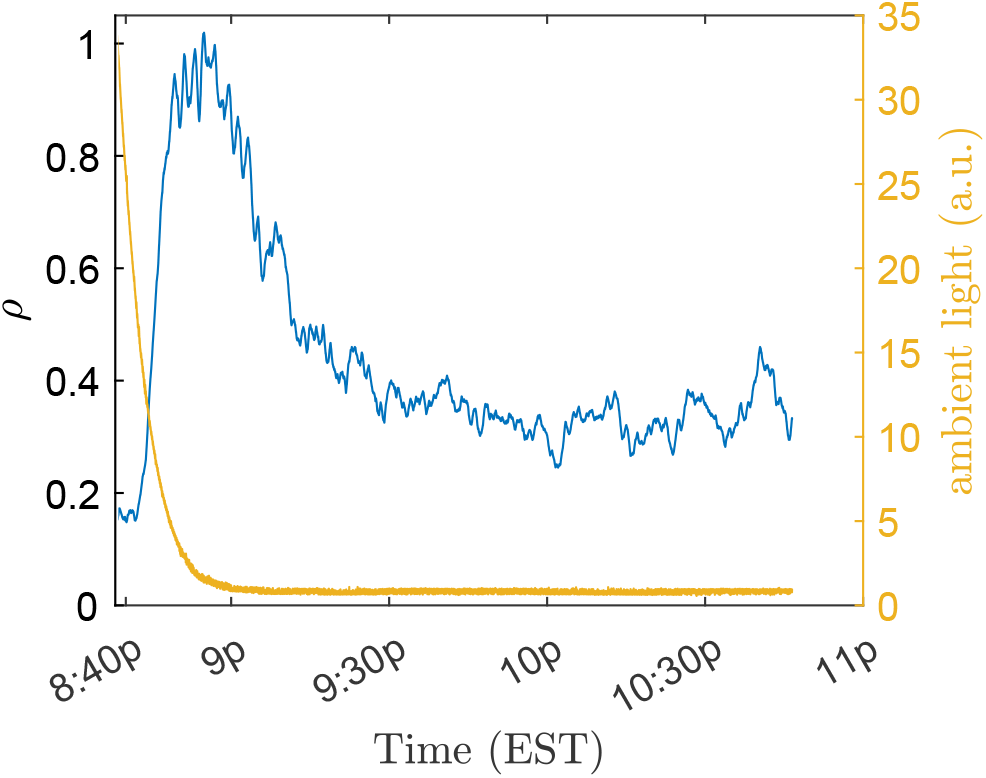
(Blue) Firefly density *ρ* calculated as the 3min moving average of the number of flashes per frame (8:38pm to 10:45pm; May 23, 2021). (Yellow) Frame brightness, quantified by average pixel intensity.

### 3 Distribution of velocities

Individual velocities are defined as the displacement between flashes divided by the corresponding interflash interval. The 3D reconstruction of the swarm shows that movement is primarily horizontal (Fig. S3A). However, the distribution of their orientations in the horizontal plane does not appear uniform. Because only a cone-shaped portion of the swarm can be reconstructed due to the camera’s limited field of view (Fig. 3B), this anisotropy could cause artifacts in the statistics of displacement. Consequently, we look at the distribution of velocities within an artificially delineated circular portion of the reconstructed swarm (Fig. 3B). We find that even these velocities show a centripetal bias (Fig. 3C), suggesting that this collective drift is real and not a mere experimental artifact. We surmise that this drift is simply due to a general tendency of fireflies to move towards the center of the lekking parcel, which is within the field of view (Fig. 1A), in order to maintain a general cohesion of the swarming structure.

**Figure S3:**
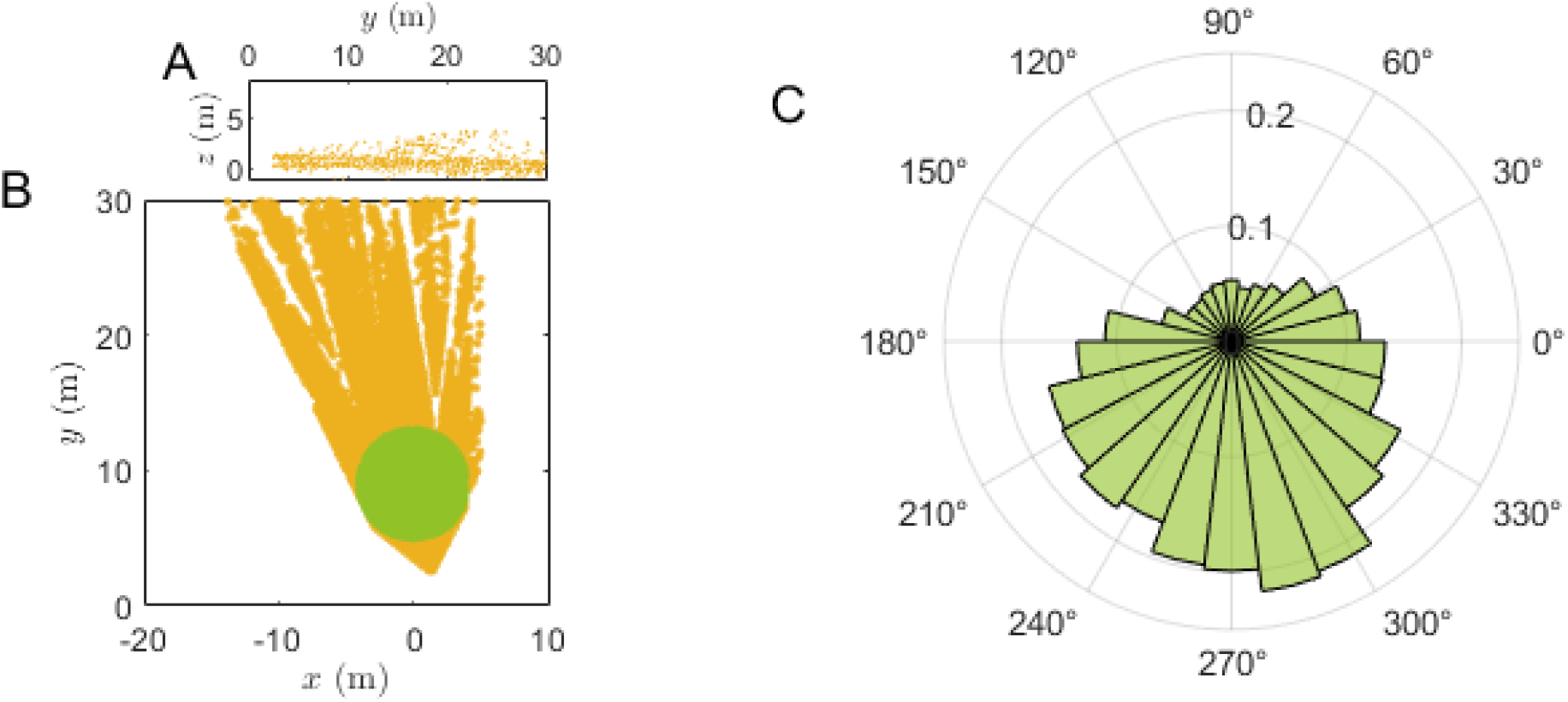
(A) Cross-section of the reconstructed swarm in the vertical plane. (B) Top-down view of the 3D swarm. The green disk is an isotropic subset of all triangulated flashes. (C) Distribution of velocity directions in the horizontal plane, corresponding to only some displacements occurring in the green disk in B.

### 4 Data

We are providing the three-dimensional reconstructions of natural swarms for May 20 to May 30, 2021 as Matlab structure files. The .xyzt field in particular provides the 3D coordinates, with xy defining the horizontal plane, z is the vertical axis (going up), and t is the corresponding time, expressed as frame number. These were recorded at 60 frames-per-second.

